# Nonconsumptive predator effects on prey demography: Dogwhelk cues decrease benthic mussel recruitment

**DOI:** 10.1101/172692

**Authors:** Sonja M. Ehlers, Ricardo A. Scrosati, Julius A. Ellrich

**Author notes:** Corresponding author: R A. Scrosati.

## Abstract

Predators have often been shown to have nonconsumptive effects (NCEs) on prey behaviour, but the demographic consequences for prey remain poorly known. This is important to understand because demography influences the impact of a species in its community. We used an intertidal predator–prey system to investigate predator NCEs on prey recruitment, a key demographic process for population persistence. Pelagic mussel larvae are known to avoid waterborne cues from dogwhelks, which prey on intertidal mussels. Through a field experiment done in Atlantic Canada, we manipulated the presence of dogwhelks in intertidal habitats during the mussel recruitment season. We measured mussel recruitment in collectors that could be reached by waterborne dogwhelk cues but not by dogwhelks themselves. We found that the nearby presence of dogwhelks significantly decreased mussel recruit density. A previous study done in the same habitats under the same experimental conditions showed that dogwhelk cues also limit the recruitment of barnacles, another prey item for dogwhelks. However, such NCEs were four times stronger than those observed for mussel recruitment. This difference relates well to the higher ability of mussels to escape predation, as mussels can relocate while barnacles cannot. Therefore, basic features of natural history may be useful to predict predator NCEs on prey recruitment.

## Introduction

Nonconsumptive effects (NCEs) of predators on prey are ubiquitous in nature. When organisms of a prey species detect cues from nearby predators, a variety of responses are often triggered to limit predation risk (Ferrari *et al*., 2010; Brönmark & Hansson, 2012). As cues from a predator can reach many prey organisms at the same time, NCEs can be extensive in prey populations (Preisser *et al*., 2005; Peacor *et al*., 2013). Thus, understanding what prey traits are affected and how has become an important research line in ecology (Weissburg *et al*., 2014).

Immediate prey responses are typically behavioural. They include moving away to minimize the chance of being reached by predators or limiting movements to avoid being detected by predators (Keppel & Scrosati, 2004; Molis *et al*., 2011; Johnston *et al*., 2012; Matassa *et al*., 2016, Johnson *et al*., 2017). The consequences of such behavioural responses for prey demography have received, however, little attention (Creel *et al*., 2007; Schoener & Spiller, 2012; Ellrich *et al.*, 2016a). This is important to understand because demography ultimately determines to a large extent the function of a species in its community. This paper focuses on predator NCEs on prey recruitment, which is a key demographic process for population persistence (Caley *et al*., 1996; Palumbi & Pinsky, 2014).

Benthic invertebrates with pelagic larvae are useful model organisms for this kind of research. For instance, a laboratory experiment has shown that larvae of blue mussels (*Mytilus edulis*) avoid waterborne chemical cues from predatory dogwhelks (*Nucella lapillus*; Morello & Yund, 2016). Dogwhelks feed on benthic mussel stages, not on their pelagic larvae (Hunt & Scheibling, 1998). However, larval avoidance of dogwhelk cues may have evolved to aid settlement-seeking larvae to find habitats with a reduced predation pressure for juveniles and adults. Such an avoidance behaviour might ultimately decrease benthic recruitment (the addition of new organisms to a benthic population after larval settlement and metamorphosis). In fact, field experiments in intertidal habitats have shown that cues from *N. lapillus* limit barnacle (*Semibalanus balanoides*) recruitment (Ellrich *et al.*, 2015a,b). This barnacle is another important prey for *N. lapillus* and it also has pelagic larvae, which settle elsewhere when dogwhelk cues are detected (Ellrich *et al.*, 2016a). Thus, the mussel–dogwhelk system offers the opportunity to start evaluating how broadly predator NCEs can limit the recruitment of benthic invertebrate prey. Through a field experiment, the present study tests the hypothesis that dogwhelk cues limit intertidal mussel recruitment.

Basic differences in natural history between mussels and barnacles may influence the intensity of such NCEs, however. The location of a barnacle is fixed for life after a larva settles and metamorphoses into a recruit (Jenkins *et al*., 2000). However, mussel recruits can detach themselves from the substrate and relocate (Bayne, 1964; Le Corre *et al*., 2013). Additionally, older mussels can immobilize dogwhelks through the production of byssus threads (Farrell & Crowe, 2007). These processes provide mussels with opportunities to escape predation that barnacles lack. Thus, we also predict that the expected dogwhelk NCEs on mussel recruitment are weaker than the NCEs recently reported for barnacles.

## Materials and Methods

We did the experiment in rocky intertidal habitats from Deming Island (45° 12′ 45″ N, 61° 10′ 26″ W), on the Atlantic coast of Nova Scotia (Canada), between May–July 2016. These habitats are constituted by stable bedrock and are protected from direct oceanic swell by rocky formations. Maximum water velocity measured with dynamometers (see design in Bell & Denny, 1994) during the study period was 6.0 ± 0.3 m s^-1^ (mean ± SE, *n* = 48). These wave-sheltered habitats were used in previous years to demonstrate that dogwhelk cues limit barnacle recruitment (Ellrich & Scrosati, 2016; Ellrich *et al.*, 2015b, 2016b). In-situ temperature measured every 30 minutes during the study period using submersible loggers (HOBO Pendant Logger, Onset Computer Corp., Pocasset, MA, USA) was 12.8 ± 0.1 °C (mean ± SE, *n* = 7 loggers). Coastal seawater salinity measured on 21 May 2016 with a refractometer was 35 ‰.

The dogwhelk used for this study was *Nucella lapillus*, which is the only dogwhelk species on the studied coast (Scrosati & Heaven, 2007). On the Atlantic coast of Nova Scotia, two blue mussel congeners, *Mytilus edulis* and *M. trossulus*, co-occur (Tam & Scrosati, 2011, 2014) and are preyed upon by *N. lapillus* (Hunt & Scheibling, 1998). These mussel species show only subtle morphological differences (Innes & Bates, 1999) and can form hybrids (Riginos & Cunningham, 2005). Thus, their visual identification is very difficult, especially at the recruit stage. Therefore, recruit counts in this study were done as *Mytilus* spp., as commonly done in ecological field studies involving these species (Cusson & Bourget, 2005; Le Corre *et al*., 2013).

We evaluated dogwhelk cue effects on mussel recruitment by manipulating dogwhelk presence in cages attached to the intertidal substrate. Each cage (Fig. 1) was made using a PVC ring (25 cm in diameter and 2.5 cm tall) and plastic mesh (0.5 cm x 0.5 cm of opening size). Each cage was divided by mesh into a central compartment (area =144 cm^2^) and a peripheral compartment (area = 347 cm^2^). The peripheral compartment was used to create two dogwhelk treatments (presence vs. absence) by enclosing either 10 dogwhelks (2.23 ± 0.02 cm in shell length, mean ± SE, *n* = 104) or none. The used dogwhelk density (ca. 3 individuals dm^-2^) was representative of the studied coast (Ellrich & Scrosati, 2016). The central compartment held a plastic mesh scourer (Our Compliments Poly Pot Scrubbers, Mississauga, ON, Canada) attached with cable ties (Fig. 1). Mesh scourers have often been used to measure intertidal mussel recruitment (Menge & Menge, 2013; South, 2016), as scourers resemble habitats where mussel larvae preferentially settle (filamentous algae or byssal threads of established mussels; Menge, 1992; Le Corre *et al*., 2013). For *Mytilus edulis* and *M. trossulus*, pelagic pediveliger larvae of at least approximately 0.25 mm in shell length settle in those habitats and, then, undergo metamorphosis, becoming recruits (Bayne, 1965; Menge *et al*., 2009; Martel *et al*., 2014). After growing to a shell length of about 0.5 mm (Hunt & Scheibling, 1996; Le Corre *et al*., 2013), such recruits may enter a second pelagic dispersal phase (Bayne, 1964). For instance, recruits of *M. edulis* up to 2.5 mm long can passively drift in the water aided by a byssus thread (Sigurdsson *et al*., 1976). In our study, observations under a stereomicroscope indicated that 70-80 % of the recruits found in the scourers belonged to the first phase, the remaining organisms belonging to the second phase. Precise counts are unavailable because the threshold size between both phases is not accurately known (Le Corre *et al*., 2013). As all of those organisms ultimately contribute to mussel recruitment (Le Corre *et al*., 2013), at the end of the experiment we counted the recruits of both phases together to determine recruit density for each scourer, as often done in field studies of this kind (Menge & Menge, 2013).

**Figure 1.**
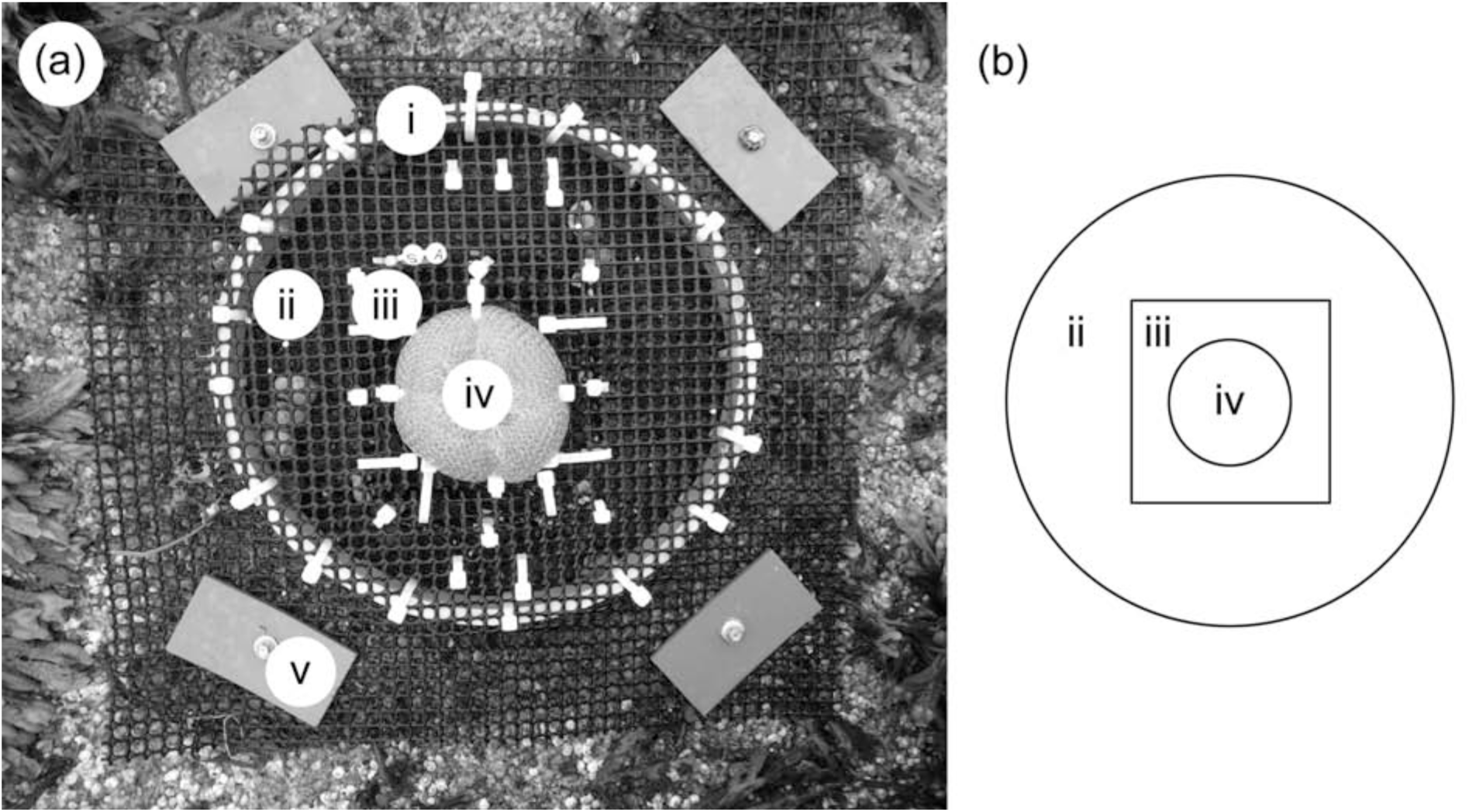
(a) Top view of a cage, showing: (i) the PVC ring determining the cage’s shape (25 cm in diameter and 2.5 cm tall), (ii) the peripheral compartment (which had 10 dogwhelks or none, depending on the treatment), (iii) the central compartment with (iv) the mesh scourer to collect mussel recruits, and (v) the four plates used to secure the cage to the intertidal substrate. (b) Simplified diagram of a cage, showing: (ii) its peripheral compartment, (iii) its central compartment, and (iv) the mesh scourer.

We set up the experiment on 21 May 2016 following a randomized complete block design with replicated treatments within blocks (Quinn & Keough, 2002). We established 12 blocks, each one including two replicate cages of each of the two dogwhelk treatments, thus yielding 24 replicates for each dogwhelk treatment. Block size was 7.7 ± 0.4 m^2^ (mean ± SE, *n* = 12 blocks) and the distance between cages within blocks was at least 0.5 m. We established the blocks at an intertidal elevation of 0.9 ± 0.1 m (mean ± SE, *n* = 12 blocks) above chart datum (the full vertical intertidal range is 1.8 m). We attached the cages to the substrate using PVC plates and screws (Fig. 1). Before installing the cages, we removed all seaweeds (mainly *Ascophyllum nodosum* and *Fucus vesiculosus*) and benthic invertebrates from the substrate to avoid chemical and physical influences from those organisms (Johnson & Strathmann, 1989; Jenkins *et al*., 1999; Beermann *et al*., 2013). During the experiment, we kept these areas devoid of free-living dogwhelks. We did not feed the caged dogwhelks during the experiment but, to prevent their starvation, we replaced them every 10-14 days with mussel-fed dogwhelks that were kept in separate cages tens of meters away from the blocks. We used mussel-fed dogwhelks because prey reacts strongly to chemical cues from predators fed conspecific prey (Cheung *et al*., 2006; Weissburg & Beauvais, 2015; Scherer & Smee, 2016). We ended the experiment on 29 July 2016, when we took all of the scourers to the laboratory to measure mussel recruit density.

In the laboratory, we stored the scourers in a freezer to preserve the integrity of the recruits until each scourer was analyzed. To count the recruits in a scourer, we unrolled the scourer and manually rinsed it in tap water to separate the recruits from the mesh. The recruits were retained in a sieve (0.212 mm × 0.212 mm of opening size) and then transferred to a Petri dish. We subsequently counted the recruits under a stereomicroscope. For each scourer, we calculated mussel recruit density by dividing the encountered number of recruits by the total area of the scourer. This standardization was necessary because small area differences could exist among the replicate scourers provided by the vendor. To calculate the total area of a scourer, we first unrolled the scourer. Then, we used scissors to cut alongside the resulting cylindrical mesh to produce a two-dimensional mesh, which we extended flat on a table. As mussel recruits occurred on both sides of this surface, we calculated the total area of the scourer as the area of that two-dimensional mesh viewed from the top multiplied by two. We evaluated the effect of dogwhelk cues (fixed factor with two levels: dogwhelk presence and absence) on mussel recruit density through an analysis of variance (ANOVA) that was appropriate for a randomized complete block design with replicated treatments within blocks (random factor with 12 levels). We confirmed the homoscedasticity and normality assumptions using Cochran’s C-test and the Kolmogorov-Smirnov test, respectively.

We also conducted a side experiment to verify that the presence of dogwhelks in a cage did not alter water motion at the place of attachment of the mesh scourer. For this purpose, we established 24 different cages on the shore on 1 June 2016. Each of those cages held a gypsum piece (Jonsson *et al*., 2006; Beermann *et al*., 2013) in the same place in which the cages used for the main experiment held a mesh scourer. We prepared the gypsum pieces following Howerton & Boyd (1992). We determined the initial dry mass of each gypsum piece to the nearest 0.1 mg. Twelve randomly selected cages each contained 10 dogwhelks in the peripheral compartment, whereas the other 12 cages lacked dogwhelks. On 2 June 2016, we collected the gypsum pieces, dried them at 60 °C for 24 h, and then measured the percent loss of mass for each piece. We compared percent loss of gypsum mass between both treatments with a t-test. We conducted all of the data analyses with STATISTICA 13.5 (Statsoft, Tulsa, OK, USA).

## Results

The ANOVA for the field experiment revealed that the presence of dogwhelks decreased intertidal mussel recruitment (Table 1). On average, mussel recruit density was 13 % lower with nearby dogwhelks than in their absence (Fig. 2). Blocks had a significant effect on mussel recruit density (Table 1), but that result merely indicates that mussel recruitment differed among blocks. The important result is that the interaction between the dogwhelks factor and the blocking factor was not significant (Table 1), indicating that the negative dogwhelk NCEs on mussel recruitment were spatially consistent on the shore. The side field experiment revealed that the presence of dogwhelks in the cages did not affect water motion (*t*_22_ = 1.14, *P* = 0.267) in the place in which the cages used for the main experiment held a mesh scourer.

**Figure 2.**
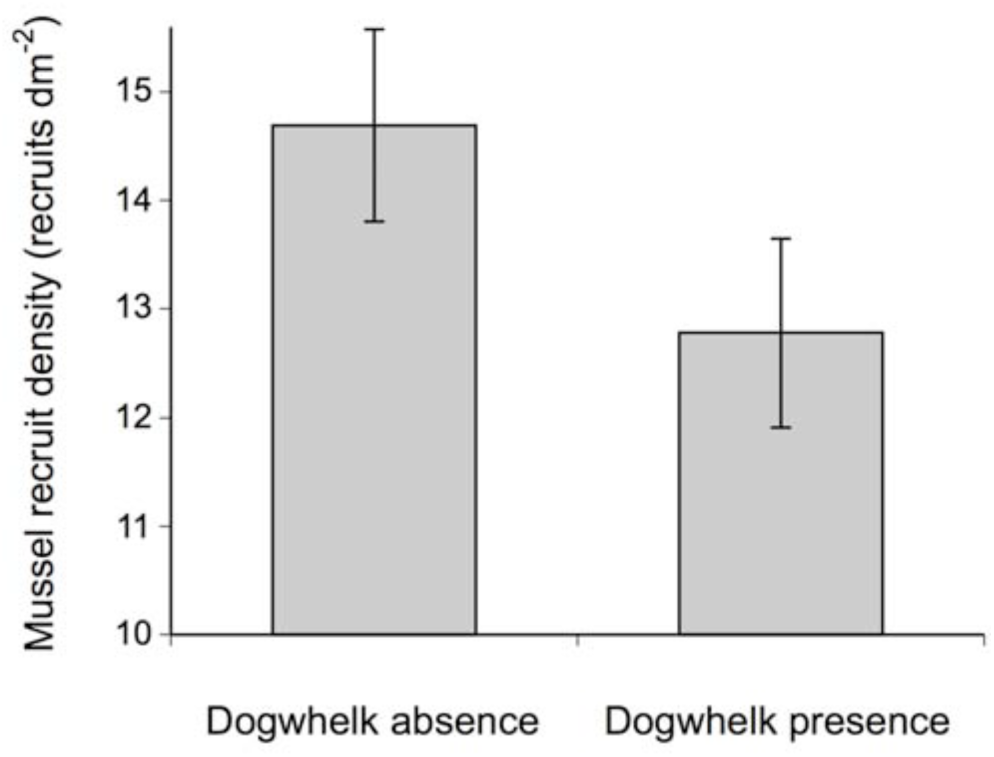
Mussel recruit density (recruits dm^-2^, mean ± SE, *n* = 24) depending on the presence or absence of dogwhelks during the field experiment.

**Table 1.**
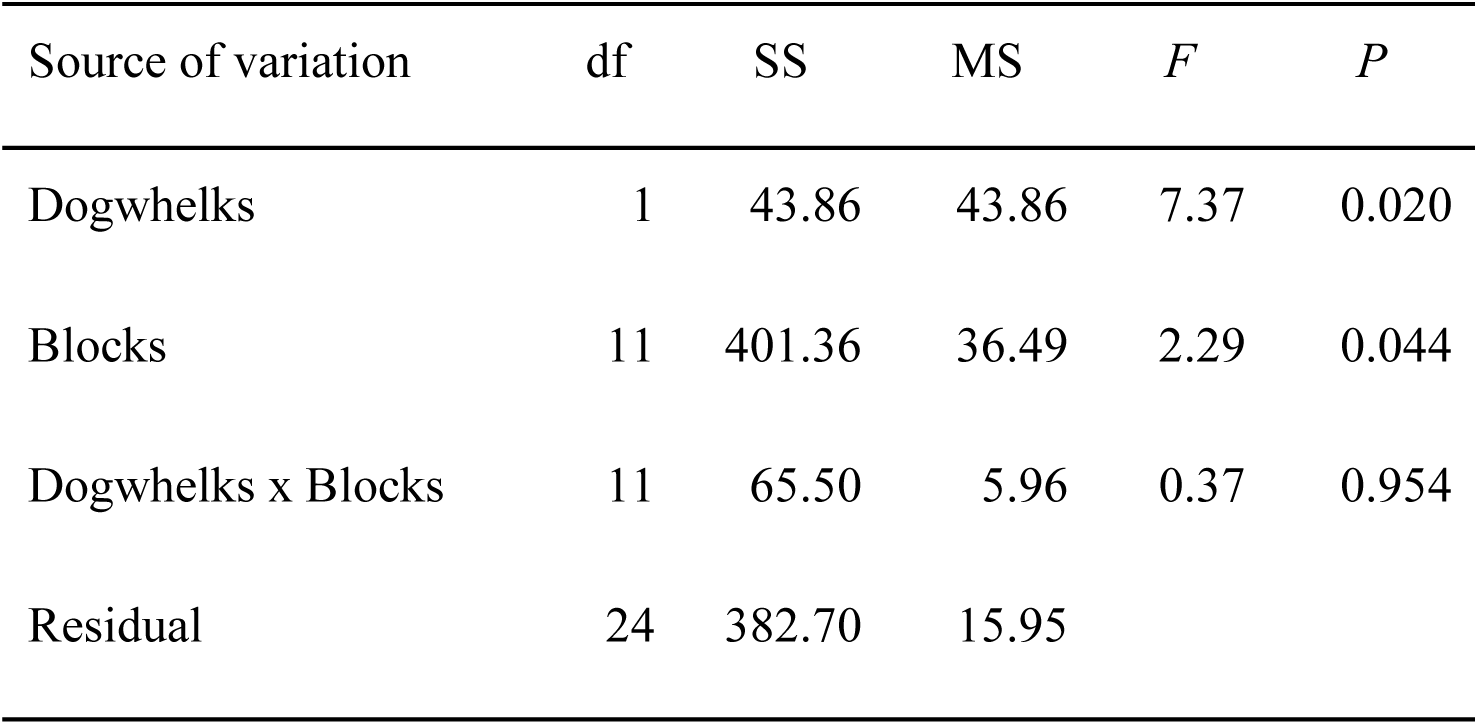
Results of the ANOVA conducted to evaluate dogwhelk NCEs on mussel recruitment.

## Discussion

This study has experimentally demonstrated that cues from predatory dogwhelks decrease mussel recruitment in intertidal habitats. This is a valuable contribution because it adds to the growing literature that is revealing predator NCEs on prey demography. Other studies have shown negative NCEs on prey reproduction (Selden *et al*., 2009; Zanette *et al*., 2011; Ellrich *et al.*, 2016a) and also recruitment (Creel *et al*., 2011; Ellrich *et al.*, 2015a; Benkwitt, 2017). These studies are important because most NCE research to date has focused on behavioural responses in prey (Ferrari *et al*., 2010; Brönmark & Hansson, 2012; Schoener & Spiller, 2012), likely because of the short times required to document such responses. Evaluating the demographic consequences requires more time, but this knowledge is necessary to better understand predator NCEs on prey population dynamics (Weissburg *et al*., 2014).

*Nucella lapillus* preys on blue mussels (Crothers, 1985). Young *N. lapillus* consume juvenile mussels and even recently hatched *N. lapillus* prey on young mussels by drilling a hole through their shells (Largen, 1967). Hence, dogwhelks are a threat to various age classes of mussels. Such an extended predation pressure is, therefore, what may have selected for the larval avoidance behaviour (Morello & Yund, 2016) that can ultimately decrease recruitment. In intertidal habitats, dogwhelks are patchily distributed (Johnson *et al*., 1998) and have a restricted activity range (Crothers, 1985; Fretter & Graham, 1994; Carro *et al*., 2012). Thus, by avoiding dogwhelk cues, young mussels likely contribute to limiting future predation risk.

This study has also revealed that the recruitment limitation caused by dogwhelk cues is weaker for mussels than for barnacles. Barnacles cannot change their location once recruited (Anderson, 1994) and cues from *Nucella lapillus* were found to limit barnacle (*Semibalanus balanoides*) recruitment by 50 % in the same habitats where we conducted the present study and under the same dogwhelk density (Ellrich *et al.*, 2016b). Mussels are also sessile, but to some extent they can relocate across the substrate throughout their benthic existence (Bayne, 1964; Hunt & Scheibling, 2002; de Vooys, 2003; van de Koppel *et al*., 2008). Older mussels can also immobilize dogwhelks using byssus threads (Farrell & Crowe, 2007). Therefore, mussels have more opportunities to escape predation than barnacles, which might explain why mussel recruitment is less responsive to dogwhelk cues than barnacle recruitment.

Pre-recruitment avoidance of predator cues has been found not only for mussels and barnacles, but also for lobsters (Boudreau *et al*., 1993), crabs (Welch *et al*., 1997; Banks & Dinnel, 2000; Tapia-Lewin & Pardo, 2014), and sea urchins (Metaxas & Burdett-Coutts, 2006). Thus, negative predator NCEs on prey recruitment might be common in benthic invertebrates with pelagic dispersal stages. The intensity of such NCEs may depend, as discussed above and among other factors, on the capacity of benthic prey stages to relocate across the substrate.

Indirect NCEs of predators on third species mediated by the direct NCEs on the predator’s prey have often been reported (Molis *et al*., 2011; Schoener & Spiller, 2012; Matassa *et al*., 2016). Those studies have generally evaluated effects on only one or a few of such third species (although exceptions evaluating responses on entire assemblages exist; Hammill *et al*., 2015). Intertidal mussels are foundation species (Altieri & van de Koppel, 2014), because they often occur in extensive patches that host several small species among the mussels (Valdivia & Thiel, 2006; O’Connor & Crowe, 2007; Arribas *et al*., 2014). Therefore, by nonconsumptively limiting mussel recruitment, dogwhelks have the potential to alter intertidal species composition. Evaluating this possibility would enrich models of community organization that currently consider only the consumptive effects of predators on foundation species and its associated biodiversity (Bruno *et al*., 2003; Scrosati *et al*., 2011).

Overall, the present study shows that predator NCEs limit intertidal mussel recruitment, potentially affecting mussel population dynamics. Moreover, the study has linked behavioural observations obtained in the laboratory (Morello & Yund, 2016) to population processes occurring under natural conditions. The field nature of our experiment is important because the complexity of intertidal environments cannot be replicated in laboratory settings. Thus, our approach agrees with recent calls to study predator NCEs under realistic conditions in order to advance NCE theory further (Weissburg *et al*., 2014; Babarro *et al*., 2016).

## Acknowledgements

We thank Jadine Krist and Zachary Sherker for field assistance. This project was funded by a Discovery Grant from the Natural Sciences and Engineering Research Council (NSERC) and a Leaders Opportunity Grant from the Canada Foundation for Innovation (CFI) awarded to R.A.S. and by a postdoctoral fellowship from the German Academic Exchange Service (DAAD) awarded to J.A.E.

## References

Altieri, A.H. & van de Koppel, J. (2014). Foundation species in marine ecosystems. In Marine community ecology and conservation: 37–56. Bertness, M.D., Bruno, J.F., Silliman, B.R. & Stachowicz, J.J. (Eds). Sunderland: Sinauer.

Anderson, D.T. (1994). Barnacles. Structure, function, development, and evolution. London: Chapman & Hall.

Arribas, L.P., Donnarumma, L., Palomo, M.G. & Scrosati, R.A. (2014). Intertidal mussels as ecosystem engineers: their associated invertebrate biodiversity under contrasting wave exposures. Mar. Biodiv. 44: 203–211.

Babarro, Vázquez E. & Olabarria, C. (2016). Importance of phenotypic plastic traits on invasive success: response of *Xenostrobus securis* to the predatory dogwhelk *Nucella lapillus*. Mar. Ecol. Prog. Ser. 560: 185–198.

Banks, J. & Dinnel, P. (2000). Settlement behavior of Dungeness crab (*Cancer magister* Dana, 1852) megalopae in the presence of the shore crab, *Hemigrapsus* (Decapoda, Brachyura). Crustaceana 73: 223–234.

Bayne, B.L. (1964). Primary and secondary settlement in *Mytilus edulis* L. (Mollusca). J. Anim. Ecol. 33: 513–523.

Bayne, B.L. (1965). Growth and the delay of metamorphosis of the larvae of *Mytilus edulis* (L.) Ophelia 2: 1-47.

Beermann, A.J., Ellrich, J.A., Molis, M. & Scrosati, R.A. (2013). Effects of seaweed canopies and adult barnacles on barnacle recruitment: the interplay of positive and negative influences. J. Exp. Mar. Biol. Ecol. 448: 162–170.

Bell, E.C. & Denny, M.W. (1994). Quantifying “wave exposure”: a simple device for recording maximum velocity and results of its use at several sites. J. Exp. Mar. Biol. Ecol. 156: 199–215.

Benkwitt, C.E. (2017). Predator effects on reef fish settlement depend on predator origin and recruit density. Ecology 98: 896–902.

Boudreau, B., Bourget, E. & Simard, Y. (1993). Behavioural responses of competent lobster postlarvae to odor plumes. Mar. Biol. 117: 63–69.

Brönmark, C. & Hansson, L.A. (2012). Chemical ecology in aquatic systems. Oxford: Oxford University Press.

Bruno, J.F., Stachowicz, J.J. & Bertness, M.D. (2003). Inclusion of facilitation into ecological theory. TrendsEcol. Evol. 18: 119–125.

Caley, M.J., Carr, M.H., Hixon, M.A., Hughes, T.P., Jones, G.P. & Menge, B.A. (1996). Recruitment and the local dynamics of open marine populations. Annu. Rev. Ecol. Syst. 27: 477–500.

Carro, B., Quintela, M., Ruiz, J.M. & Barreiro, R. (2012). AFLPs reveal different population genetic structure under contrasting environments in the marine snail *Nucella lapillus*. PLoS ONE 7: e49776.

Cheung, S.G., Luk, K.C. & Shin, P.K.S. (2006). Predator-labeling effect on byssus production in marine mussels *Perna viridis* (L.) and *Brachidontes variabilis* (Krauss). J. Chem. Ecol. 32: 1501–1512.

Creel, A., Christianson, D., Liley, S. & Winney, J.A. (2007). Predation risk affects reproductive physiology and demography of elk. Science 315: 960.

Creel, A., Christianson, D. & Winney, J.A. (2011). A survey of the effects of wolf predation risk on pregnancy rates and calf recruitment in elk. Ecol. Appl. 21: 2847–2853.

Crothers, J.H. (1985). Dog-whelks: an introduction to the biology of *Nucella lapillus* (L.). Field Studies 6: 291–360.

Cusson, M. & Bourget, E. (2005). Small-scale variations in mussel (*Mytilus* spp.) dynamics and local production. J. Sea Res. 53: 255–268.

de Vooys, C.G.N (2003). Effect of a tripeptide on the aggregational behaviour of the blue mussel *Mytilus edulis*. Mar. Biol. 142: 1119–1123.

Ellrich, J.A. & Scrosati, R.A. (2016). Water motion modulates predator nonconsumptive limitation of prey recruitment. Ecosphere 7: e01402.

Ellrich, J.A., Scrosati, R.A., Bertolini, C. & Molis, M. (2016a). A predator has nonconsumptive effects on different life-history stages of a prey. Mar. Biol. 163: article 5.

Ellrich, J.A., Scrosati, R.A. & Molis, M. (2015a). Predator nonconsumptive effects on prey recruitment weaken with recruit density. Ecology 96: 611–616.

Ellrich, J.A., Scrosati, R.A. & Petzold, W. (2015b). Predator density affects nonconsumptive predator limitation of prey recruitment: field experimental evidence. J. Exp. Mar. Biol. Ecol. 472: 72–76.

Ellrich, J.A., Scrosati, R.A., Romoth, K. & Molis, M. (2016b). Adult prey neutralizes predator nonconsumptive limitation of prey recruitment. PLoS ONE 11: e0154572.

Farrell, E.D. & Crowe, T.P. (2007). The use of byssus threads by *Mytilus edulis* as an active defense against *Nucella lapillus*. J. Mar. Biol. Assoc. U. K. 87: 559–564.

Ferrari, M.C.O. Wisenden, B.D. & Chivers, D.P. (2010). Chemical ecology of predator-prey interactions in aquatic ecosystems: a review and prospectus. Can. J. Zool. 88: 698–724.

Fretter, V. & Graham, A. (1994). British prosobranch molluscs. Their functional anatomy and ecology. London: The Ray Society.

Hammill, E., Atwood, T.B. & Srivastava, D.S. (2015). Predation threat alters composition and functioning of bromeliad ecosystems. Ecosystems 18: 857–866.

Howerton, R.D. & Boyd, C.E. (1992). Measurement of water circulation in ponds with gypsum blocks. Aquacult. Eng. 11: 141–155.

Hunt, H.L. & Scheibling, R.E. (1996). Physical and biological factors influencing mussel (*Mytilus trossulus, M. edulis*) settlement on a wave-exposed rocky shore. Mar. Ecol. Prog. Ser. 142: 135–145.

Hunt, H.L. & Scheibling, R.E. (1998). Effects of whelk (*Nucella lapillus* (L.)) predation on mussel (*Mytilus trossulus* (Gould), *M. edulis* (L.)) assemblages in tidepools and on emergent rock on a wave-exposed rocky shore in Nova Scotia, Canada. J. Exp. Mar. Biol. Ecol. 226: 87–113.

Hunt, H.L. & Scheibling, R.E. (2002). Movement and wave dislodgment of mussels on a wave-exposed rocky shore. Veliger 45: 273–277.

Innes, D.J. & Bates, J.A. (1999). Morphological variation of *Mytilus edulis* and *Mytilus trossulus* in eastern Newfoundland. Mar. Biol. 133: 691–699.

Jenkins, S.R., Åberg, P., Cervin, G., Coleman, R.A., Delany, J., Della Santina, P., Hawkins, S.J., LaCroix, E., Myers, A.A., Lindegarth, M., Power, A.M., Roberts, M.F. & Hartnoll, R.G. (2000). Spatial and temporal variation in settlement and recruitment of the intertidal barnacle *Semibalanus balanoides* (L.) (Crustacea: Cirripedia) over a European scale. J. Exp. Mar. Biol. Ecol. 243: 209–225.

Jenkins, S.R., Norton, T.A. & Hawkins, S.J. (1999). Settlement and post-settlement interactions between *Semibalanus balanoides* (L.) (Crustacea: Cirripedia) and three species of fucoid canopy algae. J. Exp. Mar. Biol. Ecol. 236: 49–67.

Johnson, G.C., Karajah, M.T., Mayo, K., Armenta, T.C. & Blumstein, D.T. (2017). The bigger they are the better they taste: size predicts predation risk and anti-predator behavior in giant clams. J. Zool. 301: 102–107.

Johnson, L.E. & Strathmann, R.R. (1989). Settling barnacle larvae avoid substrata previously occupied by a mobile predator. J. Exp. Mar. Biol. Ecol. 128: 7–103.

Johnson, M.P., Hughes, R.N., Burrows, M.T. & Hawkins, S.J. (1998). Beyond the predation halo: small scale gradients in barnacle populations affected by the relative refuge value of crevices. J. Exp. Mar. Biol. Ecol. 231: 163–170.

Johnston, B.R., Molis, M. & Scrosati, R.A. (2012). Predator chemical cues affect prey feeding activity differently in juveniles and adults. Can. J. Zool. 90: 128–132.

Jonsson, P.R., Granhag, L., Moschella, P.S., Åberg, P., Hawkins, S.J. & Thompson, R.C. (2006). Interactions between wave action and grazing control the distribution of intertidal macroalgae. Ecology 87: 1169–1178.

Keppel, E. & Scrosati, R. (2004). Chemically mediated avoidance of *Hemigrapsus nudus* (Crustacea) by *Littorina scutulata* (Gastropoda): effects of species coexistence and variable cues. Anim. Behav. 68: 915–920.

Largen, M.J. (1967). The diet of the dog-whelk, *Nucella lapillus* (Gastropoda Prosobranchia). J. Zool. 151: 123–127.

Le Corre, N., Martel, A.L., Guichard, F. & Johnson, L.E. (2013). Variation in recruitment: differentiating the roles of primary and secondary settlement of blue mussels *Mytilus* spp. Mar. Ecol. Prog. Ser. 481: 133–146.

Martel, A.L., Tremblay, R., Toupoint, N., Olivier, F. & Myrand, B. (2014). Veliger size at metamorphosis and temporal variability in prodissoconch II morphometry in the blue mussel (*Mytilus edulis*): potential impact on recruitment. J. Shellfish Res. 33: 443–455.

Matassa, C.M., Donelan, S.C., Luttbeg, B. & Trussell, G.C. (2016). Resource levels and prey state influence antipredator behavior and the strength of nonconsumptive predator effects. Oikos 125:1478–1488.

Menge, B.A. (1992). Community regulation: under what conditions are bottom-up factors important on rocky shores?. Ecology 73: 755–765.

Menge, B.A., Chan, F., Nielsen, K.J., Di Lorenzo, E. & Lubchenco, J. (2009). Climatic variation alters supply-side ecology: impact of climate patterns on phytoplankton and mussel recruitment. Ecol. Monogr. 79: 379–395.

Menge, B.A. & Menge, D.N.L. (2013). Dynamics of coastal meta-ecosystems: the intermittent upwelling hypothesis and a test in rocky intertidal regions. Ecol. Monogr. 83: 283–310.

Metaxas, A. & Burdett-Coutts, V. (2006). Response of invertebrate larvae to the presence of the ctenophore *Bolinopsis infundibulum*, a potential predator. J. Exp. Mar. Biol. Ecol. 334: 187–195.

Molis, M., Preuss, I., Firmenich, A. & Ellrich, J. (2011). Predation risk indirectly enhances survival of seaweed recruits but not intraspecific competition in an intermediate herbivore species. J. Ecol. 99: 807–817.

Morello, S.L. & Yund, P.O. (2016). Response of competent blue mussel (*Mytilus edulis*) larvae to positive and negative settlement cues. J. Exp. Mar. Biol. Ecol. 480: 8–16.

O’Connor, N.E. & Crowe, T.P. (2007). Biodiversity among mussels: separating the influence of sizes of mussels from the ages of patches. J. Mar. Biol. Assoc. U. K. 87: 551–557.

Palumbi, S.R. & Pinsky, M.L. (2014). Marine dispersal, ecology, and conservation. In Marine community ecology and conservation: 57–83. Bertness, M.D., Bruno, J.F., Silliman, B.R. & Stachowicz, J.J. (Eds). Sunderland: Sinauer.

Peacor, S.D., Peckarsky, B.L., Trussell, G.C. & Vonesh, J.R. (2013). Costs of predator-induced phenotypic plasticity: a graphical model for predicting the contribution of nonconsumptive and consumptive effects of predators on prey. Oecologia 171: 1–10.

Preisser, E.L., Bolnick, D.I. & Benard, M.F. (2005). Scared to death? The effects of intimidation and consumption in predator–prey interactions. Ecology 86: 501–509.

Quinn, G.P. & Keough, M.J. (2002). Experimental design and data analysis for biologists. Cambridge: Cambridge University Press.

Riginos, C. & Cunningham, C.W. (2005). Local adaptation and species segregation in two mussel (*Mytilus edulis x Mytilus trossulus*) hybrid zones. Mol. Ecol. 14: 381–400.

Scherer, A.E. & Smee, D.L. (2016). A review of predator diet effects on prey defensive responses. Chemoecology 26: 83–100.

Schoener, T.W. & Spiller, D.A. (2012). Perspective: kinds of trait-mediated indirect effects in ecological communities. A synthesis. In Trait-mediated indirect interactions: ecological and evolutionary perspectives: 9–27. Ohgushi, T., Schmitz, O.J. & Holt, R.D. (Eds) Cambridge: Cambridge University Press.

Scrosati, R. & Heaven, C. (2007). Spatial trends in community richness, diversity, and evenness across rocky intertidal environmental stress gradients in eastern Canada. Mar. Ecol. Prog. Ser. 342: 1–14.

Scrosati, R.A., van Genne, B., Heaven, C.S. & Watt, C.A. (2011). Species richness and diversity in different functional groups across environmental stress gradients: a model for marine rocky shores. Ecography 34: 151–161.

Selden, R., Johnson, A.S. & Ellers, O. (2009). Waterborne cues from crabs induce thicker skeletons, smaller gonads, and size-specific changes in growth rate in sea urchins. Mar. Biol. 156:1057–1071.

Sigurdsson, J.B., Titman, C.W. & Davies, P.A. (1976). The dispersal of young post-larval bivalve molluscs by byssus threads. Nature 262: 386–387.

South, P.M. (2016). An experimental assessment of measures of mussel settlement: effects of temporal, procedural, and spatial variations. J. Exp. Mar. Biol. Ecol. 482: 64–74.

Tam, J.C. & Scrosati, R.A. (2011). Mussel and dogwhelk distribution along the north-west Atlantic coast: testing predictions derived from the abundant-centre model. J. Biogeogr. 38: 1536–1545.

Tam, J.C. & Scrosati, R.A. (2014). Distribution of cryptic mussel species (*Mytilus edulis* and*M. trossulus*) along wave exposure gradients on northwest Atlantic rocky shores. Mar. Biol. Res. 10: 51–60.

Tapia-Lewin, S. & Pardo, L.M. (2014). Field assessment of the predation risk–food availability trade-off in crab megalopae settlement. PLoS ONE 9: e95335.

Valdivia, N. & Thiel, M. (2006). Effects of point-source nutrient addition and mussel removal on epibiotic assemblages in *Perumytilus purpuratus* beds. J. Sea Res. 56: 271–283.

van de Koppel, J., Gascoigne, J.C., Theraulaz, G., Rietkerk, M., Mooij, W.M. & Herman, P.M.J. (2008). Experimental evidence for spatial self-organization and its emergent effects in mussel bed ecosystems. Science 322: 739–742.

Weissburg, M. & Beauvais, J. (2015). The smell of success: the amount of prey consumed by predators determines the strength and range of cascading non-consumptive effects. PeerJ 3: e1426.

Weissburg, M., Smee, D.L. & Ferner, M.C. (2014). The sensory ecology of nonconsumptive predator effects. Am. Nat. 184: 141–157.

Welch, J.M., Rittschof, D., Bullock, T.M. & Fordward, R.B. (1997). Effects of chemical cues on settlement behaviour of blue crab *Callinectes sapidus* postlarvae. Mar. Ecol. Prog. Ser. 154: 143–153.

Zanette, L.Y., White, A.F., Allen, M.C. & Clinchy, M. (2011). Perceived predation risk reduces the number of offspring songbirds produce per year. Science 334: 1398–1401.

